# Discovering gene regulatory networks of multiple phenotypic groups using dynamic Bayesian networks

**DOI:** 10.1101/2021.12.16.473035

**Authors:** Polina Suter, Jack Kuipers, Niko Beerenwinkel

**Affiliations:** Department of Biosystems Science and Engineering, ETH Zurich, Matternstrasse 26, 4058, Basel, Switzerland; SIB Swiss Institute of Bioinformatics, Switzerland

**Keywords:** dynamic Bayesian networks, time series, gene expression, Bayesian learning, classification, MCMC

## Abstract

Dynamic Bayesian networks (DBNs) can be used for the discovery of gene regulatory networks from time series gene expression data. Here, we suggest a strategy for learning DBNs from gene expression data by employing a Bayesian approach that is scalable to large networks and is targeted at learning models with high predictive accuracy. Our framework can be used to learn DBNs for multiple groups of samples and highlight differences and similarities in their gene regulatory networks. We learn these DBN models based on different structural and parametric assumptions and select the optimal model based on the cross-validated predictive accuracy. We show in simulation studies that our approach is better equipped to prevent overfitting than techniques used in previous studies. We applied the proposed DBN-based classification approach to two time series transcriptomic datasets from the Gene Expression Omnibus database, each comprising data from distinct phenotypic groups of the same tissue type. In the first case, we used DBNs to characterize responders and non-responders to anti-cancer therapy. In the second case, we compared normal to tumor cells of colorectal tissue. The classification accuracy reached by the DBN-based classifier for both datasets was higher than reported previously. For the colorectal cancer dataset, our analysis suggested that GRNs for cancer and normal tissues have a lot of differences, which are most pronounced in the neighborhoods of oncogenes and known cancer tissue markers. The identified differences in gene networks of cancer and normal cells may be used for the discovery of targeted therapies.

## 1. Introduction

Learning gene regulatory networks (GRNs) from gene expression data has been the focus of much research in the last decades [1, 2, 3]. The precise knowledge of GRNs can help to understand the molecular mechanisms driving diseases and facilitate the search for targeted therapies [4, 5]. More recently, a lot of research was focused on learning gene-gene interaction networks that are specific to certain contexts, for example, particular tissues or disease subtypes [6, 7, 8]. Moreover, it has been shown experimentally that different mutations of the same gene can cause distinct changes in signaling pathways [9]. These studies suggest that gene interactions differ between different phenotypic groups of the same tissue or disease type. Moreover, context-specific GRNs can facilitate the discovery of targeted therapies [10]. However, phenotypic differences are mostly ignored in studies devoted to learning GRNs *de novo*.

Multiple computational methods can be used to learn GRNs from observational data, including correlation analysis [11], Boolean networks [12], Bayesian networks [13, 14], and differential equation models [15, 16]. Bayesian network approaches provide a good trade-off between the scalability and interpretability of discovered networks but do not allow directed cycles, rendering it impossible for them to model feedback loops. The Dynamic Bayesian Network (DBN) model overcomes this problem by including dependencies between nodes at different time points and accommodating the possibility of cycles [17]. DBNs were previously used for inference of biological networks [18], including GRNs [19, 20, 21, 22, 23] and multi-omics networks [24]. However, none of these studies considered non-homogeneous datasets where DBN structures may differ between groups of samples. Kourou et al. [25] were the first to employ DBNs for classification, however, their method was not scalable to network structures beyond 40 nodes.

Structure learning of DBNs is a computationally challenging problem. Existing approaches are either not scalable [25, 23, 26], use greedy search [20] or restrict the DBN topology to decrease computational costs and accommodate larger networks [18, 24, 26, 27]. However, any restriction may potentially result in the discovery of suboptimal models [28].

The goal of this study is to create a scalable framework for learning DBN models that provide high predictive accuracy and can be used for learning GRNs for multiple subgroups of samples, defined, for example, by molecular, histological, or clinical phenotypes. We employed a Bayesian approach [29] for learning DBNs, which is scalable to networks with hundreds of nodes and implemented in the R-package BiDAG [30]. Our DBN learning strategy provides means to prevent overfitting of the DBN structure by filtering out low-confidence edges and has not been applied to learning large GRNs before. We extended the functionality of BiDAG such that it can be used for classification and learning DBNs with parameters that change over time.

Previous studies demonstrated better performance of the Bayesian approach to structure learning compared to non-Bayesian approaches [29, 8, 31]. However, these simulations did not compare consensus models obtained via sampling from the posterior and bootstrapping in the case of greedy hill-climbing. In addition, limiting the parent set size was used as a way to prevent overfitting in DBNs [18, 24] and to decrease computational costs [26, 27]. However, the impact of uniform restrictions of the parent set sizes on the structure fit and predictive accuracy in large networks in the high dimensional setting was never explored. In simulation studies, we demonstrated how the Bayesian approach to structure learning of DBNs is better equipped to prevent overfitting compared to greedy hill-climbing coupled with a hard limit on the number of parents per node. Among other ways to improve predictive accuracy, we propose investigating models that use prior biological information about associations between genes or prohibit edges between nodes within the same time slice.

To demonstrate the advantages of the described approach, we identified time-series datasets in the Gene Expression Omnibus (GEO) database, that included gene expression data for at least two different phenotypic groups of the same tissue and comprised at least 50 observations in each of two consecutive time slices. We found two datasets (GSE5462 and GSE37182) that satisfied these criteria. The main question we wanted to answer using our framework was if the phenotypic groups in each dataset could be better represented by DBNs with the same structure (but not parameters) or if gene regulation differs substantially and the groups are more accurately represented by different DBN structures. The results of our analysis aligned well with previously validated biological findings, while the DBN-based classifier demonstrated a higher classification accuracy of patient samples with regard to phenotypic groups than was reported in [25] for both datasets.

## 2. Methods and data

A DBN is a probabilistic graphical model for the joint distribution of random variables **X** = (*X*_1_,…, *X_n_*) observed at time points *t* = 0, 1,…, *T*. The DBN model uses a directed graph to encode a factorization of the joint distribution of (**X**^*t*^) along the time slices *t* = 0,…, *T* (Fig. 1A). Here we consider DBNs in which structures are identical for all time slices. We also assume that variables in time slice *t* can depend on other variables in time slice *t* and on variables in time slice *t* − 1, i.e.

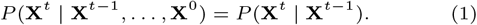

**Fig. 1.**
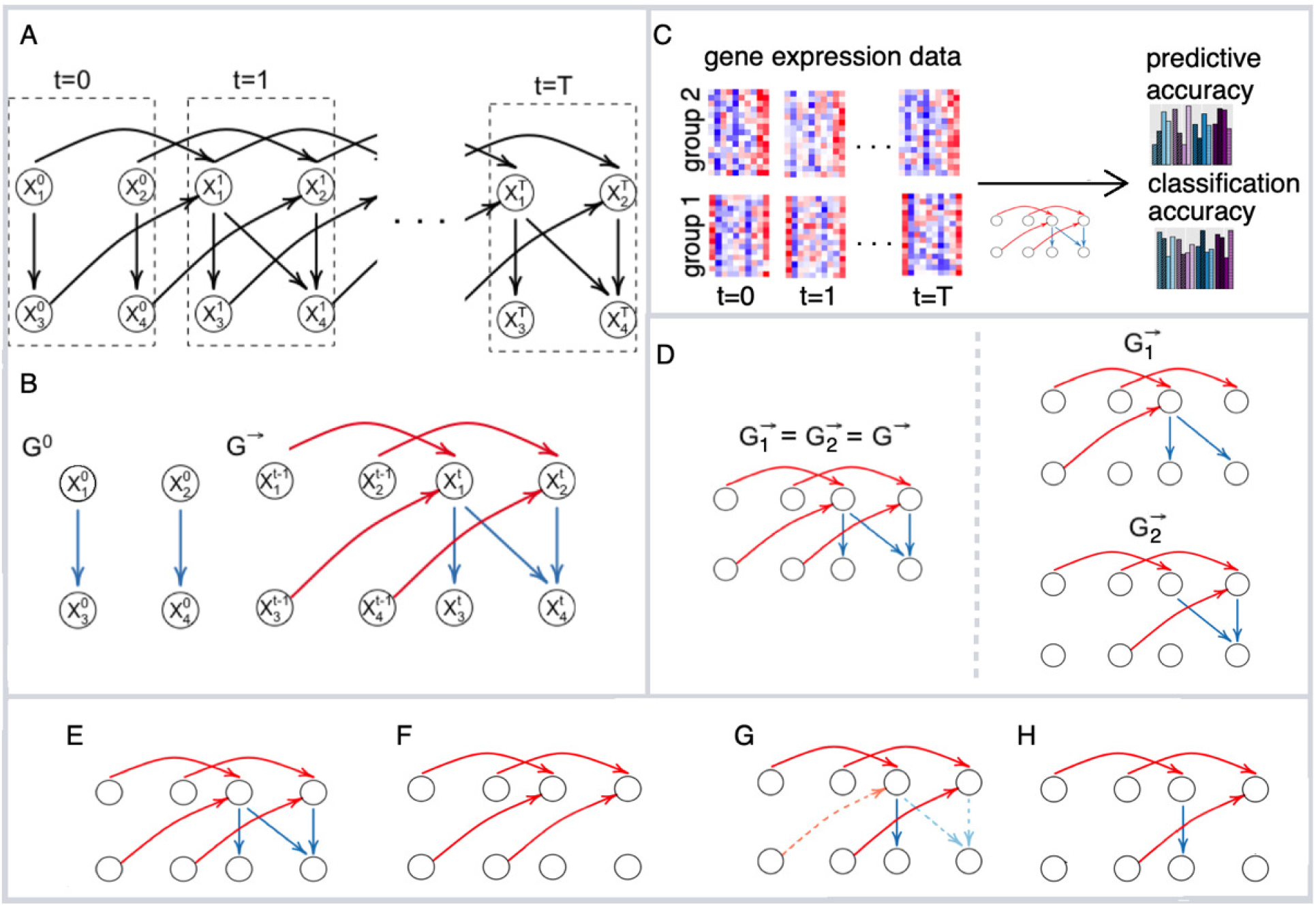
DBN learning and classification framework. (A) The unfolded structure of the first-order DBN model consisting of *T* + 1 time slices can be represented by initial and transition structures. (B) The edges between time slices are highlighted in red and called inter-edges. The edges within time slices are highlighted in blue and called intra-edges. (C) We learn DBN-based classification models from the time-series gene expression data of two phenotypic groups (group 1 and group 2) with various structural and parametric assumptions and assess the predictive accuracy and classification accuracy of these models using leave-one-out cross-validation.(D) We evaluate models where phenotypic groups are represented by the same or different DBN structures. For each model we consider a set of four structural restrictions/prior assumptions: (E) no restrictions, (F) model without intra-edges (G) model that penalizes non-STRING edges (dashed) (H) model where non-STRING edges are blacklisted.

Such DBN models are referred to as first-order DBNs.

The joint probability distribution of a DBN with *T* + 1 time slices is

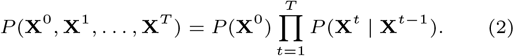

With these assumptions, the unfolded structure of a DBN (Fig. 1A) can be represented in a compact way with two directed acyclic graphs (*G*^0^, *G*^→^) which are referred to as initial structure and transition structure, respectively (Fig. 1B). The edges within one time slice are called intra-edges and edges between time slices are called inter-edges.

Within each time slice *t* > 0 the joint distribution of *X*_1_,…, *X_n_* is factorized according to a Bayesian network model:

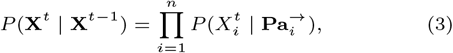

where 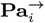 denotes the set of parents of node 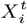 in time slices *t* and *t* − 1 in *G*^→^. For *G*^0^ the parent sets 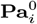 are used instead to factorize *P*(**X**^0^).

To fully specify a DBN, we also need parameters *θ* which describe probabilistic dependencies between each node 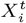 and its parents in a DBN structure. We assume that 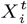 are jointly normally distributed. This results in the distribution of each node 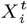 being a linear regression on its parents [32]:

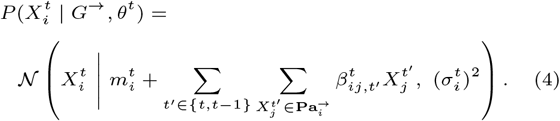

For each time slice *t*, we have the parameters *θ^t^* = (*m^t^, B^t^*, (*σ*^2^)^*t*^), where *m^t^* is a vector of regression intercepts, *B^t^* = (*β_ij,t′_*)^*t*^ a set of all regression coefficients and (*σ*^2^)^*t*^ a vector of variances. For *G*_0_, the sum over parents in the previous time slice is dropped. We consider two cases, namely stationary DBNs where *θ*^1^ = … = *θ^T^* ≕ *θ*^→^ and non-stationary DBNs, where *θ*^1^,…, *θ^T^* are generally different. The parameters *θ*^0^ and *θ*^→^ are different even for a stationary model. In a non-stationary model, we assume non-stationarity of parameters, while the structure *G*^→^ is assumed to be the same across time slices 1,…, *T*.

We also consider a special case where the initial structure *G*^0^ is the same as the internal structure of the transition structure *G*^→^, i.e., for all nodes, all intra-slice parents in *G*^→^ are the same as these nodes’ parents in *G*^0^.

For learning the DBN structure from observational data *D*, we employ the Bayesian approach implemented in the R package BiDAG [29, 30], which uses the BGe score for learning and sampling the structures of Bayesian networks [32, 33]. The BGe score of a graph is its posterior probability and factorizes into terms 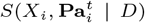, each of which depends only on a single node and its parents. For DBNs, the dataset *D* consists of *N* observations from *T* + 1 time slices. To estimate a non-stationary DBN, we divide *D* in *T* + 1 parts and define the BGe score of a DBN structure as

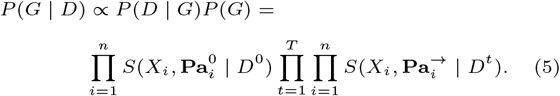

To perform structure learning for a stationary model we divide the data into 2 parts: *D*^0^ and *D*^→^, where *D*^→^ contains observations from all pairs of neighboring time slices. Equation 5 then simplifies to

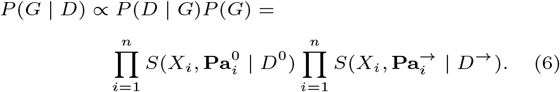

We use the iterative order Markov chain Monte Carlo (MCMC) scheme [29] to estimate the *a posteriori* (MAP) structures *G*^0^ and *G*^→^. In addition, we estimate consensus structures by averaging over a sample of graphs from the posterior distribution and composing consensus structures of edges whose posterior probability is higher that 0.9 [29, 30].

### 2.1. Learning DBN models for phenotypic subgroups

We propose a novel framework for learning the DBN models for *K* phenotypic subgroups 1,…, *K*, from time-series gene expression datasets *D*_1_, *D*_2_,…, *D_K_*. We analyzed two datasets, each comprising gene expression from *K* = 2 subgroups (Figure 1C). We propose considering two models: one which assumes that DBN structures are subgroup-specific and the another one which represents all subgroups by a single DBN structure (Figure 1D). In the latter case the differences between subgroups can be explained by differences in parameters. From a biological perspective, it is interesting to understand to which extent the interaction networks of different subgroups, for example defined by different phenotypes, differ from each other.

When dealing with gene expression data it is important to account for the fact that only a limited number of observations is usually available, which often results in poor structure fit and low predictive accuracy [31, 26]. For this reason, we suggest learning a range of models based on different assumptions regarding network structure and parameters and likewise select the model for downstream analysis based on cross-validated predictive accuracy. We use mean absolute error (MAE) as a measure of predictive accuracy, which was already used in previous DBN application studies [24, 18]. To assess how well different DBN models describe the underlying dynamic process, we predicted the values of all genes in all times slices using the true values of expression of genes at time point *t* = 0 using leave-one-out CV. We used the learned structures and parameters to predict the values of nodes in time slices consecutively using the model specified in Equation (4). Finally, we computed the MAE for each node and each time slice and averaged it across all genes, slices, and test samples. The models with lower MAE provide a higher predictive accuracy.

It is know that including prior biological knowledge can improve network learning [34], so we consider two different ways to include such knowledge: by penalizing the edges that can not be found in public protein-protein interaction databases, such as, e.g., the STRING database [35] (Figure 1g) and by excluding these edges completely from the search space (Figure 1h). Penalization is implemented by imposing a non-uniform prior over structures:

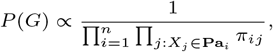

where *π_ij_* = 1 if the interaction between genes *X_i_* and *X_j_* can be found in the STRING database with a confidence level of at least 0.4, and *π_ij_* = 2 otherwise.

Many studies prohibit intra-edges, which usually makes sense when the time intervals between observations are short. We do not assume the presence or absence of intra-edges by default. Instead, we include the model without intra-edges in the set of investigated models (Figure 1F) and compare its predictive accuracy to other models, including the model without any structural restrictions (Figure 1E).

### 2.2. Classification accuracy

In addition to the predictive accuracy, we measured crossvalidated classification accuracy to evaluate how well the DBN-based classifier can discriminate between the analyzed subgroups.

For each DBN dataset, we either learned two DBN structures *G*_1_ and *G*_2_ or learned one DBN structure *G* but estimated the maximum MAP parameters separately for groups 1 and 2. After learning the networks, we estimated the MAP parameters and scored the test sample against each model by computing the likelihoods:

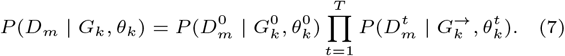

We further computed posterior probabilities of class memberships *Z_m_* of observations *D_m_* using the class prior *P*(*k*) = *τ_k_* estimated from the training data:

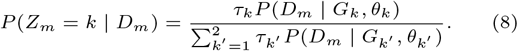

The samples were assigned to a class corresponding to the highest posterior probability. We compared assignments of test samples in all CV runs to their true assignments. For benchmarking, we compared the DBN-based classifier against both random forest and naive Bayes classifiers [36, 37]. For the random forest classifier, we ran the CV 100 times to average out randomness in the results.

### 2.3. Data

We applied the described framework to two biological datasets, each containing time-series gene expression data of two phenotypic groups of the same tissue type (Section 4).

The dataset GSE5462 contains gene expression data of 116 biopsies from 58 breast cancer patients at two time points: pre-treatment and 10-14 days after treatment with letrozole. We log2-transformed and normalized the raw data using robust multiarray averaging (RMA, R-package affy, [38]) for subsequent DBN analysis.

The second dataset, GSE37182, contains expression data of 172 biopsies from 15 colorectal cancer patients, totaling 88 normal tissue biopsies and 84 tumor tissue biopsies. The samples were obtained during surgery and left at room temperature at four time points: 20 minutes (*t* = 0), 60 minutes (*t* = 1), 180 minutes (*t* = 2), and 360 minutes (*t* = 3). Afterwards, the samples were stored at −80°C until RNA extraction. The data from the repository was already normalized separately within each group (tumor and nontumor). To make samples between two conditions comparable, we used the package NormalyzerDE [39] and performed median normalization.

### 2.4. Variable selection

To select nodes to be included in the DBNs we performed DGE analysis using the R package limma [40]. We considered genes as differentially expressed between conditions if their FDR-adjusted p-value was smaller than 0.05. In general, we did not apply a log2-fold-change cut-off, unless specified otherwise.

### 2.5. Simulation studies

We generated 50 two-step DBNs structures. For each DBN structure, we generated 30 training samples from 4 consecutive time slices and 2 test samples. We learned MAP and consensus structures corresponding to posterior thresholds of *p* ∈ {0.3, 0.5, 0.7, 0.9, 0.99} using the Bayesian approach (functions iterativeMCMC and orderMCMC from the R-package BiDAG). We also learned minimum BIC structures using greedy hill-climbing with the limits on the number of parents of 3 and 5. For each limit, we also learned consensus structures based on bootstrap support levels of *ρ* ∈ {0.3, 0.5, 0.7, 0.9, 0.99}.

To assess the predictive accuracy we estimated MAP parameters for each model and predicted the values consecutively in time slices using values in the first slice of test samples as input. We divided the resulting MAE for each model by the MAE of the ground truth structure to make results between runs comparable. We compared the learned structures to the ground truth using true positive rate (TPR), false discovery rate (FDR), and structural Hamming distance (SHD, introduced in [41]).

## 3. Results

### 3.1. Simulated data

In the simulation studies (Section 2.5, Data and code availability), we explored the situation when the number of observations between neighboring time points is smaller than the number of nodes. Maximum score structures obtained by all algorithms resulted in a high FDR (Figure 2A). However, this problem was more pronounced for the hill-climbing approach, and the FDR is higher for the structures learned using a higher limit per parent set size. As a result, the SHD between estimated maximum score structures with a limit of 3 parents per node and the ground truth structures was smaller than SHD between structures with 5 parents limit (Figure 2B). At the same time, the TPR was higher for the limit of 5 nodes than for 3 nodes. For the MCMC scheme, we did not limit the number of parents and obtained better results for the MAP structure in terms of TPR, FDR, and SHD. However, MAP structures still contained a lot of false-positive edges. Therefore, we suggest improving the structure fit by filtering out low-confidence edges. We observed that consensus structures yielded much lower SHDs than maximum scoring structures (Figure 2B). More importantly, the bootstrap-based structures for a limit of 5 parents provide a better fit than using the limit of 3 parents. The lowest SHD was reached for the MCMC scheme and a posterior level of 0.99, showing another advantage of using MCMC over greedy hill climbing.

**Fig. 2.**
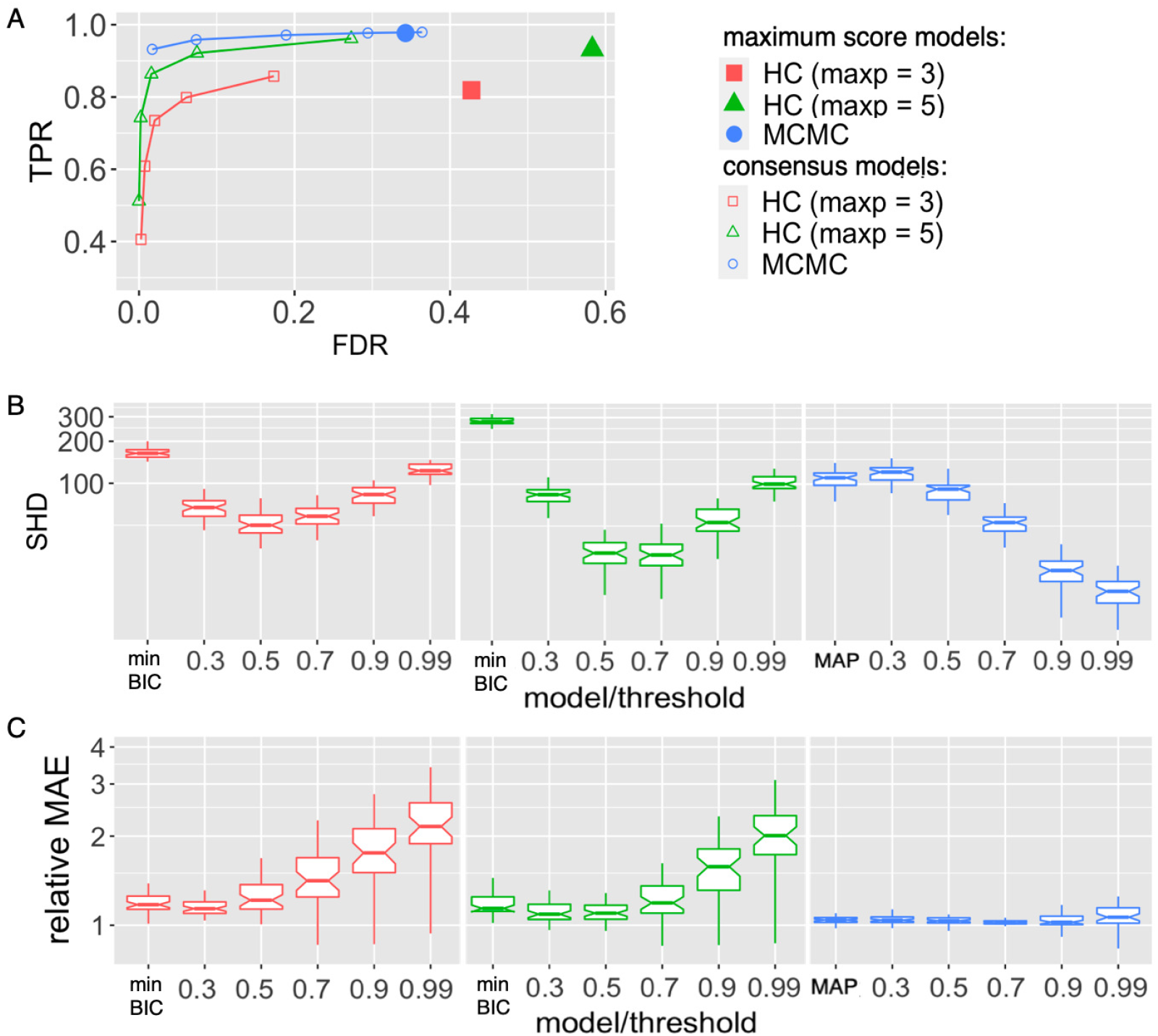
MAE and structure fit on simulated data. 50 random 2-step DBN structures were generated with *n* = 120 nodes and three parents on average for each node in the transition structure. The training datasets contained 30 samples from four consecutive time slices, the test datasets included 2 samples each. MCMC (blue) and hill-climbing (HC, red and green) algorithms were used to learn the DBN structures and compare them to the ground truth using the TPR and FDR (A) and SHD (B). The performance of the hill-climbing was evaluated for two limits for the parent set size: *maxp* = 3 (red) and *maxp* = 5 (green). Consensus models for MCMC were estimated using a range of posterior thresholds (0.3, 0.5, 0.7, 0.9, 0.99). For hill climbing, consensus models were estimated using the same range for bootstrap support levels. MAE was estimated based on 2 test samples for each simulation run and model and divided by the MAE of the ground truth structure to get comparable MAE levels between the runs (C).

For the hill-climbing approach, the relative MAE of the bootstrap-based models with the lowest SHD to ground truth structure was higher than the relative MAE of the maximum score model (Figure 2C). This finding suggests that structural overfitting does not necessarily result in a worse predictive accuracy. A possible explanation can be that false-positive edges in high-scoring structures directly connect the nodes which are indirectly connected in the ground truth graph. Such false-positive edges do not affect the MAE negatively. However, the discovery of such edges is still undesirable, and hence sparser structures should be preferred in cases when the relative MAE of several models are similar. MCMC reached the lowest mean relative MAE level of 1.08 with the posterior threshold of 0.7. For the hill-climbing, the lowest relative MAE of 1.37 was reached for the bootstrap-based structure with a support threshold of 0.3 and the limit of 5 for the maximum parent set size.

### 3.2. Analysis of time-series gene expression data

We applied the proposed approach to two transcriptomic datasets from the GEO repository (Section 2.3, Section 4): the colorectal cancer dataset GSE37182 and the breast cancer dataset GSE5462. For each dataset, we learned several DBN models (Section 2.1) using the Bayesian approach and measured, via leave-one-out cross-validation (CV), how they perform with regard to predictive accuracy and classification accuracy.

For the colorectal cancer dataset, we have learned a non-stationary DBN and compared it to a stationary DBN by computing MAE. A non-stationary DBN can describe the underlying process with higher precision. However, it can also lead to overfitting.

In addition to different structural and parametric assumptions, for each dataset, we explored DBN models containing a different number of genes. Smaller gene sets that are differentially expressed between two phenotypic groups can provide a better separation with regard to classification while larger gene sets can be more interesting for the downstream analysis and understanding differences in regulatory networks of different phenotypic groups.

### 3.3. Analysis of breast cancer time-series gene expression data

The GSE5462 dataset contains gene expression measurements for two groups of breast cancer patients: responders and nonresponders to treatment (Section 2.3). We used three sets of genes to assess how feature selection affects DBN predictive and classification accuracy. In the first gene set, we included all genes that were differentially expressed in responders compared to non-responders (Section 2.4). For the second set of genes, we identified differentially expressed genes in post-treatment samples using pre-treatment samples as reference. Finally, in the third set, we included all genes from the first set as well as genes from the second set whose absolute log2-fold-change were larger than 0.5. In addition, we included all transcription factors of the identified genes found in the database Omnipath [42]. The resulting sets of genes, denoted DBN_19, DBN_158 and DBN_125, contained 19, 158, and 125 genes, respectively.

The classification accuracy ranged from 67% to 88.4% over all DBN models (Fig. 3). The highest classification accuracy was reached with the smallest number of nodes, and the accuracy dropped with an increasing number of nodes. This finding can be explained by the variable selection process for the three sets of nodes. The smallest set of nodes includes all genes which are differentially expressed between responders and non-responders. Thus, even without any network component, it provides a strong signal for classification. The naive Bayes approach provides similarly high accuracy for this set of genes, while the random forest classifier is slightly less accurate. The largest set of genes, DBN_158, comprises all genes which are differentially expressed between time points but none of those which are differentially expressed between the subgroups. For this reason, it is not surprising that the classification accuracy is lower than for DBN_19. At the same time, in the DBN_158 and DBN_125 gene sets, the best DBN models outperform the naive Bayes classifier, which does not account for the network component.

**Fig. 3.**
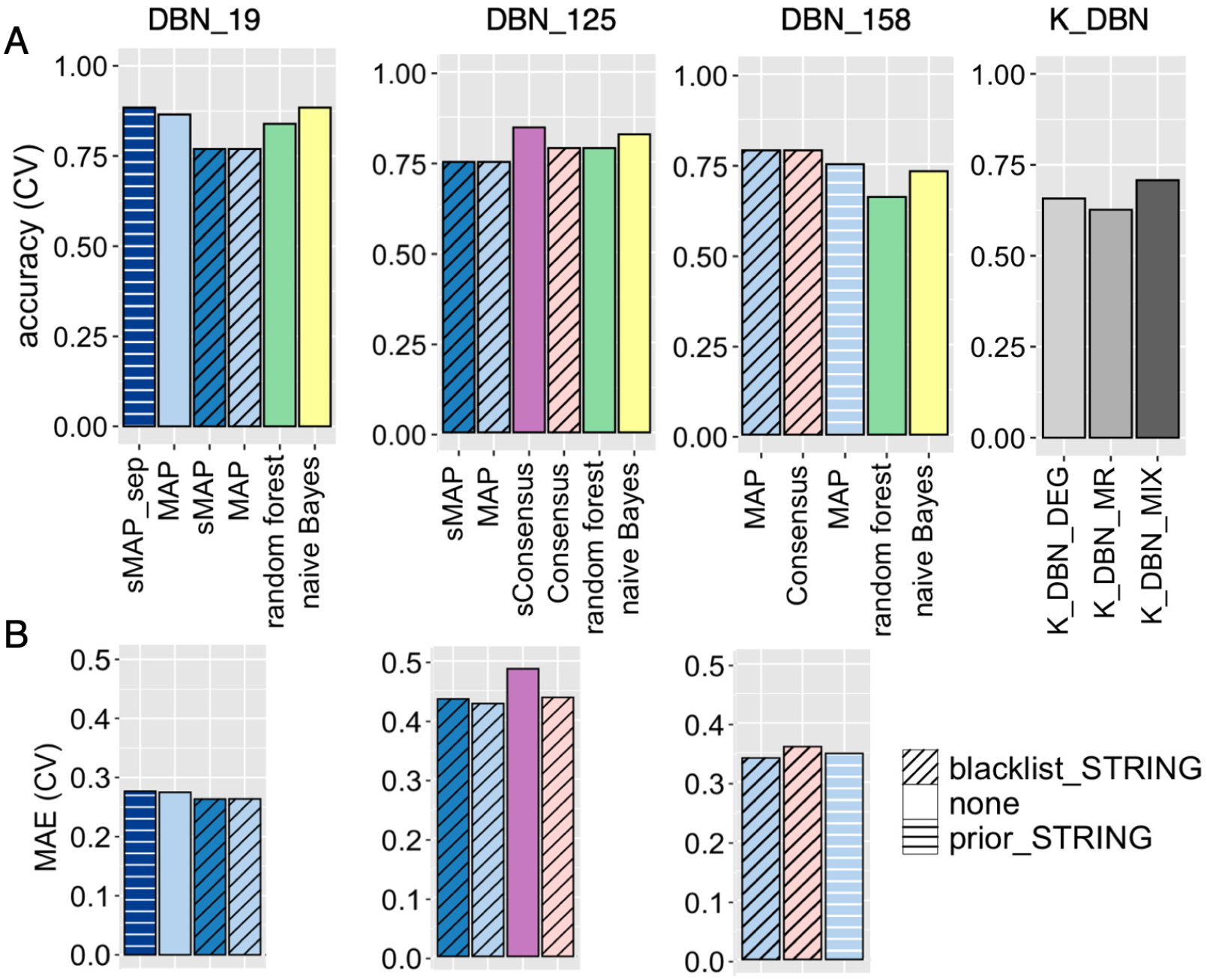
DBN CV results for the breast cancer dataset. DBN models were learned from time-series gene expression data from biopsies of responders and non-responders to the treatment. For each set of genes (DBN_19, DBN_125 and DBN_158) four best performing DBN models are shown (two with the highest accuracy and two with the lowest MAE) and the corresponding accuracy of other classifiers on the same set of genes (A). The fill color corresponds to the model used for classification: MAP and consensus DBNs assuming the same structure for responders and non-responders (MAP and Consensus, light-blue and light-pink), MAP and consensus DBN assuming different structures for responders and non-responders, and same internal structures of initial and transition structures (sMAP and sConsensus, blue and violet), MAP DBN assuming different structures for responders and non-responders and different internal structures of initial and transition structures (sMAP_sep, dark blue), random forest (green), naive Bayes classifier (yellow). For comparison, K_DBN category corresponds to leave-one-out classification accuracy reported for DBN models in [25]: DEG denotes differentially expressed genes, MR their master regulators, and MIX the union of DEG and MR. (B) Cross-validated MAE of DBN models learned in the proposed framework.

We further note that, for all gene sets, the lowest MAE was reached for DBNs learning the same DBN structure for both subgroups (Fig. 3B). This finding aligns well with the differential gene expression (DGE) and pathway enrichment analysis. Since out of 22,283 genes, only 19 were differentially expressed, we can assume that the GRNs are very similar in responders and non-responders. However, the highest classification accuracy of 88.4% was reported for models that learned DBN structures independently for responders and nonresponders for DBN_19 and DBN_125. The best models in this work outperformed the highest classification accuracy of DBN models reported in [25], which was 70.77%.

We summarized the performance of all DBN models in a global ranking (Supplementary information Table S1), which sums ranks in each accuracy measure and set of genes included in the models (Ranking of DBN models for breast cancer dataset. To identify which DBN model performs consistently better than others for networks of different sizes and incorporate both accuracy and MAE we created a ranking of models model with regard to classification accuracy and MAE for each DBN size; ties were resolved by taking maximum.Ranks of models in for each category are reported in this table.). MAP and consensus models which blacklisted non-STRING edges and learned one structure for responders and nonresponders performed best overall. Based on this ranking, for the downstream analysis, we chose the MAP model that learned the same DBN structure for responders and non-responders, included differentially expressed genes and their transcription factors and blacklisted all non-STRING interactions. Even though the classification accuracy for this model was not the highest, the lower MAE suggests that it better predicts the changes in post-treatment gene expression levels and hence is more appropriate for the downstream analysis.

Pathway enrichment analysis showed that no KEGG [43] pathway was enriched in the differentially expressed genes. However, when we assessed the set of all parent nodes of these genes in the estimated DBN structure (Figure S2), three KEGG pathways (p53 signaling, cellular senescence, and cell cycle) were enriched (FDR< 0.05). Thus, the DBN model connected genes found to be important for treatment response to genes from major cancer-related pathways. Among these genes, the most connected node was *CDK1* (Cyclin Dependent Kinase 1), which is a known target for treating breast cancer [44]. Interestingly, Cdk inhibitors are already approved for treating breast cancer as the first-line treatment in combination with letrozole (used in the analyzed dataset) [45] which confirms the discovered link.

### 3.4. Analysis of colorectal cancer time-series gene expression data

For the colorectal cancer dataset GSE37182, we performed the DGE analysis at three consecutive time points, using t=0 as a reference. The number of differentially expressed genes increased with time. In total, we identified 58 genes that were differentially expressed over all time points in both subgroups. We included them in the first set of genes DBN_58.

We proceeded with the identification of transcription factors that may be involved in regulating the identified genes using the Omnipath database. We combined them with the first set of genes and used their union (DB_122) to learn the extended DBN models. We learned multiple DBN models using various structural and parametric assumptions and performed crossvalidation as described above (Section 2.1).

The CV classification accuracy was 100% for all models and higher than the accuracy of the best model reported in [25] (98.5%). The MAE was clearly the lowest for non-stationary DBNs (Fig. 4). Among the non-stationary models, the lowest MAE was reached for models where intra-edges were blacklisted. From a biological perspective, the non-stationary model is also plausible. First, the time lags between the measurements were non-uniform. Second, the tissue was left at room temperature, and the process of degradation likely led to changes in the strengths of dependencies between genes.

**Fig. 4.**
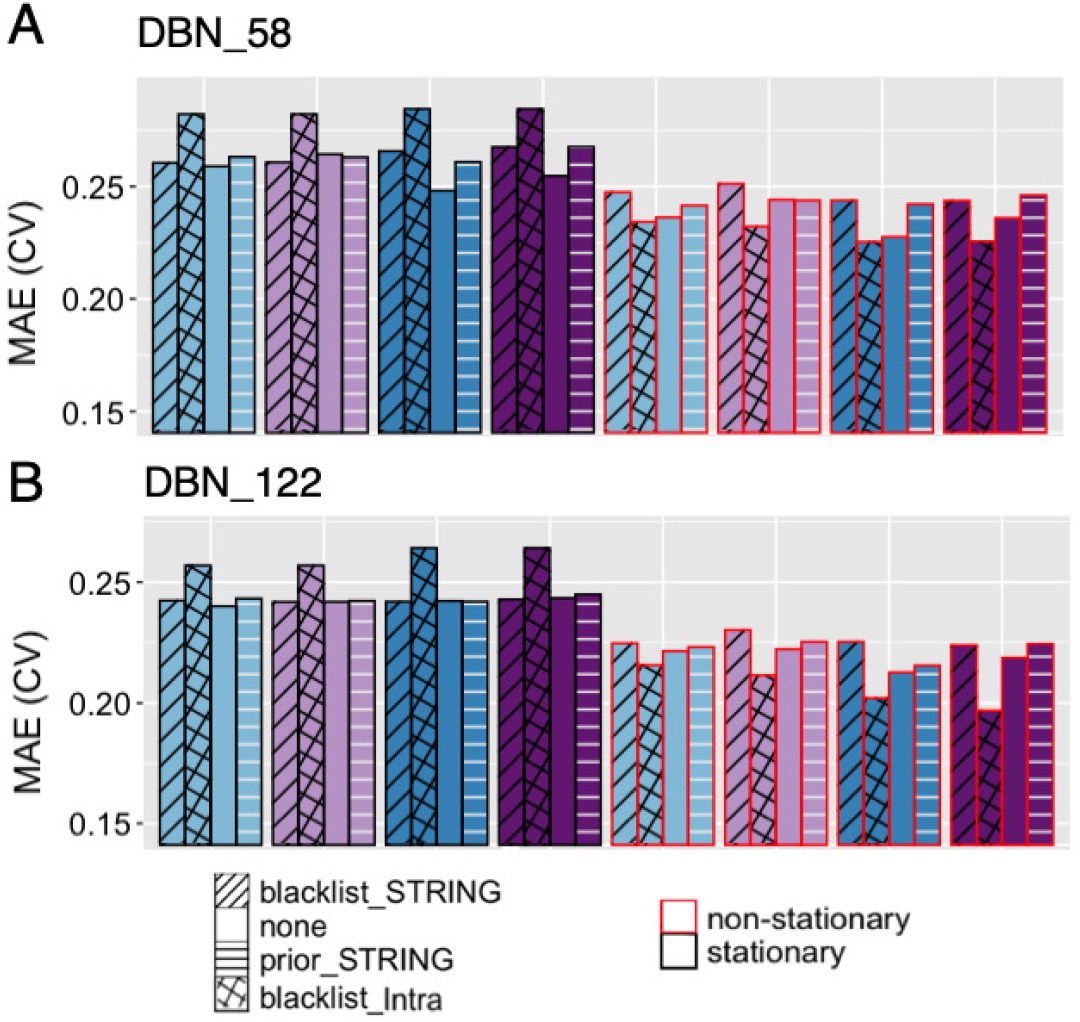
Cross-validated MAE of DBN models for GSE37182 dataset. DBN models were learned from the time-series gene expression data of normal and cancer cells from the colon. Colors correspond to MAP (blue) and consensus (violet) DBN models. Shades correspond to the similarity of DBN structures estimated for two tumor and normal samples: lighter shades represent models assuming the same DBN structure for both subgroups (MAP, Consensus), darker shades represent models assuming different DBN structures for two subgroups (sMAP, sConsensus). Patterns correspond to prior and structural constraints: STRING-based penalization (horizontal lines), STRING-based blacklisting (slanted lines), blacklisting of intra-slice edges (hatch pattern); absence of pattern corresponds to models with unrestricted topology. (A) MAE(CV) of DBNs with 58 nodes (differentially expressed genes). (B) MAE(CV) of DBNs with 122 nodes (differentially expressed genes and their transcription factors).

We observed that DBN models that learn structures for tumor and normal subgroups independently resulted in the lowest MAE. Consequently, for the downstream analysis, we selected a consensus non-stationary model which learns the DBNs for the set of genes DBN_122 separately for normal and tumor samples and blacklists intra-slice edges. The DBN models for cancer and normal subgroups shared 60% of edges. Such a high overlap suggests that a lot of underlying processes in cancer and normal cells can be described by the same dependencies between genes.

To highlight the differences and similarities between analyzed phenotypic groups, we identified the nodes with the most different and similar interaction partners in networks representing tumor and normal subgroups. There were 18 nodes that had neighborhoods with empty intersections in two networks. Out of these, three genes (*FOS, JUN, GADD45B*) belong to the KEGG colorectal cancer pathway. Two genes from this set, namely *FOSB* and *JUN*, were identified and validated as markers for colorectal tumor tissue degradation [47]. Out of 20 nodes with most similar neighborhoods 10 can be found of a generic transcription pathway (Fig 5, [46]).

**Fig. 5.**
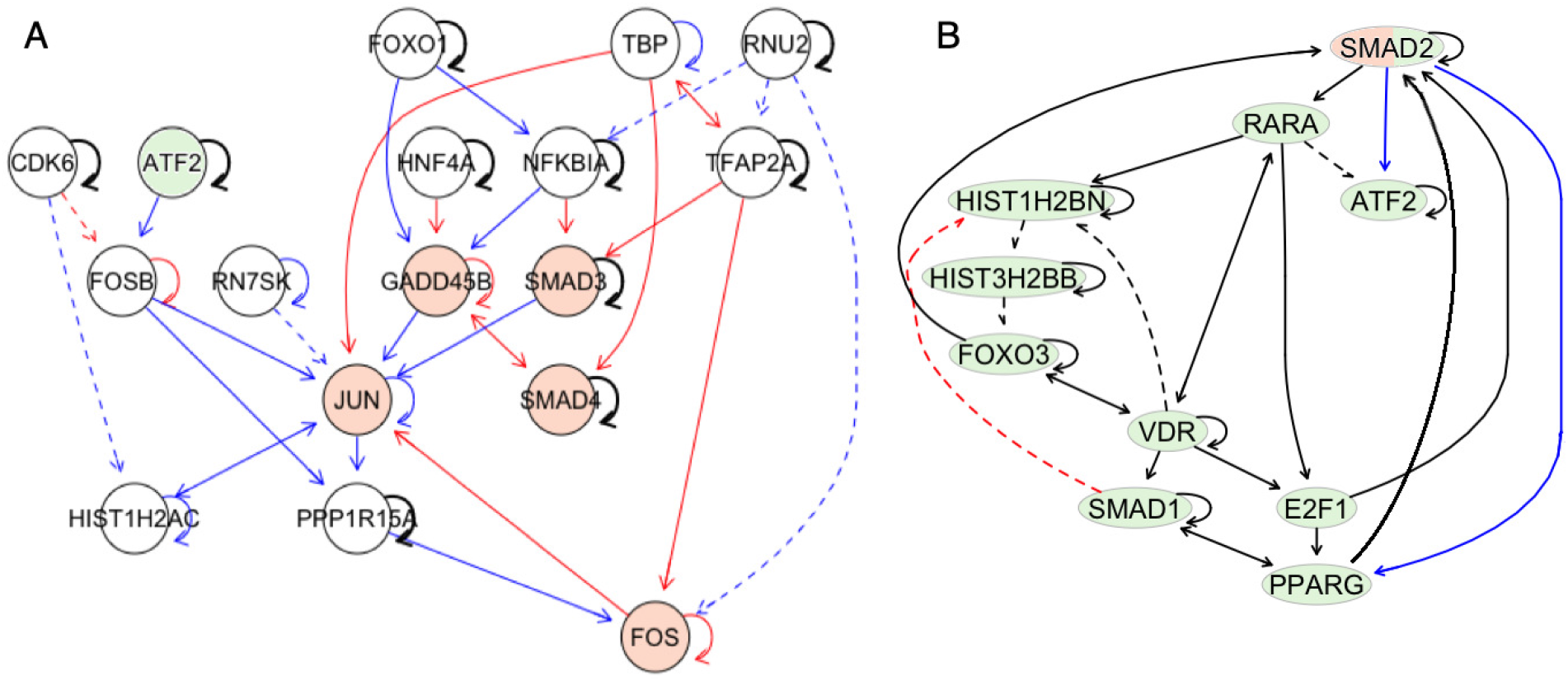
Subnetworks of DBN transition structures discovered for the GSE37182 dataset. Structures of non-stationary DBN models without intra-edges were learned for normal and tumor biopsies. Blue edges denote the edges which are specific to normal DBN only; red edges are specific to cancer DBN. Black edges are present in both models. Solid lines correspond to edges between genes which were found as interactors in the STRING database. Genes from the colorectal cancer pathway (KEGG) are highlighted in orange. (A) Most differently connected genes (*FOSB, JUN, FOS, GADD45B*) in DBN transition structures of cancer and normal DBN models that are either enriched in the colorectal cancer signaling pathway or were previously validated as biomarkers of cancer tissue for the dataset GSE37182 and their parents in the learned DBN models. (B) Most similarly connected genes, which were also found on the generic transcription pathway [46] (highlighted in green).

### 3.5. Discussion

DBNs are powerful models for analyzing time-series gene expression data because they allow us to shed light on GRNs that orchestrate molecular processes. Lately, a lot of research focused on learning context-specific gene networks. In this work, we proposed a framework for learning DBNs for multiple phenotypic groups. This framework employs the Bayesian approach to structure learning of DBNs and suggests several sets of structural and parametric assumptions to find the model with the highest predictive accuracy. We assessed predictive accuracy using cross-validation as a way to prevent overfitting which is a major problem in the analysis of high-dimensional and noisy biological data. We showed in simulation studies how the Bayesian approach allows to prevent overfitting and increase predictive accuracy compared to the hill-climbing approach.

We employed the proposed framework to analyze two time-series gene expression datasets, each comprising data from two subgroups of samples. The GSE5462 dataset included gene-expression data of breast tumor biopsies taken before and after treatment with letrozole. Our analysis suggested that gene regulatory networks do not differ substantially between responders and non-responders. The parents of the differentially expressed genes in the learned DBN structures included genes from the cell cycle and tp-53 pathways, with kinase *CDK1* being the most connected node. This situation indicates that differences in the signaling pathways of responders and non-responders might lie at the phosphoproteome level since the activity of kinases generally can not be detected from gene expression data. However, even with a few differences detected at the gene expression level, the classification accuracy we obtained was higher than reported in the previous study [25].

For the GSE37182 colon cancer dataset, different DBN structures for tumor and normal samples resulted in the lowest MAE. However, the corresponding DBN structures overlapped by 60%, and the biggest differences between networks were identified in the neighborhoods of genes from the colorectal cancer pathway as well as genes, which were previously validated as markers stratifying cancer and normal tissue. Our findings correspond to the common understanding that many housekeeping pathways work similarly in tumor and normal cells, and the biggest differences can be observed in the expression and interactions of oncogenes. In addition, the non-stationary DBN model implemented as a part of the proposed framework resulted in the lowest MAE.

In this work, we learned the DBN models of known phenotypic groups, so we were able to evaluate the classification accuracy and flag overfitting. Our results suggest that the DBN model can be useful for learning context-specific regulatory networks. At the next step, the DBN-based mixture model can be employed for the discovery of unknown disease subtypes based on time series transcriptomic or multi-omic. data.

## 4. Data and code availability

The unprocessed data is available at the public GEO repository under identifiers GSE5462 and GSE37182. The datasets can be accessed at https://www.ncbi.nlm.nih.gov/geo/query/acc.cgi?acc=GSE5462 https://www.ncbi.nlm.nih.gov/geo/query/acc.cgi?acc=GSE37182

The reproducible code and the results are available at the GitHub repository https://github.com/cbg-ethz/DBNclass.

The latest version of the BiDAG package including implemented updates is available at CRAN repository https://cran.r-project.org/web/packages/BiDAG

## 6. Acknowledgments

The authors thank the anonymous reviewers for their valuable suggestions. Part of this research was supported by the European Research Council (ERC) Synergy Grant 609883 and SystemsX.ch Research, Technology and Development (RTD) Grant 2013/150.

## 7. Author contributions statement

P.S. conceived the research project. N.B. supervised the research project. P.S. designed and implemented the computational framework and conducted the analyses. P.S. and N.B. wrote the manuscript. N.B. and J.K. reviewed and edited the manuscript.

## 8. Key points

- The proposed strategy to learn GRNs for multiple phenotypic groups unifies the efficient approach to DBN structure learning and the versatile method for model selection, enabling the discovery of models with high predictive and classification accuracy.
- The efficient Bayesian approach to structure learning is better equipped to prevent overfitting than greedy hillclimbing coupled with other conventional techniques.
- The proposed method highlights the importance of using cross-validation for model selection and assessing the degree of differences between the regulatory networks of different phenotypic groups.

**Polina Suter.** She is PhD candidate at ETH Zurich. Her research interests include context-specific network learning, Bayesian methods for network reconstruction, multi-omics data integration.

**Jack Kuipers.** He is a senior scientist at ETH Zurich. His research is focused on cancer evolution modelling, phylogenetic tree inference, probabilistic graphical models and single-cell sequencing analysis.

**Niko Beerenwinkel.** He is a professor at ETH Zurich. His research is at the interface of mathematics, statistics, and computer science with biology and medicine.

## 9. Supplementary information

**Fig. S1.**
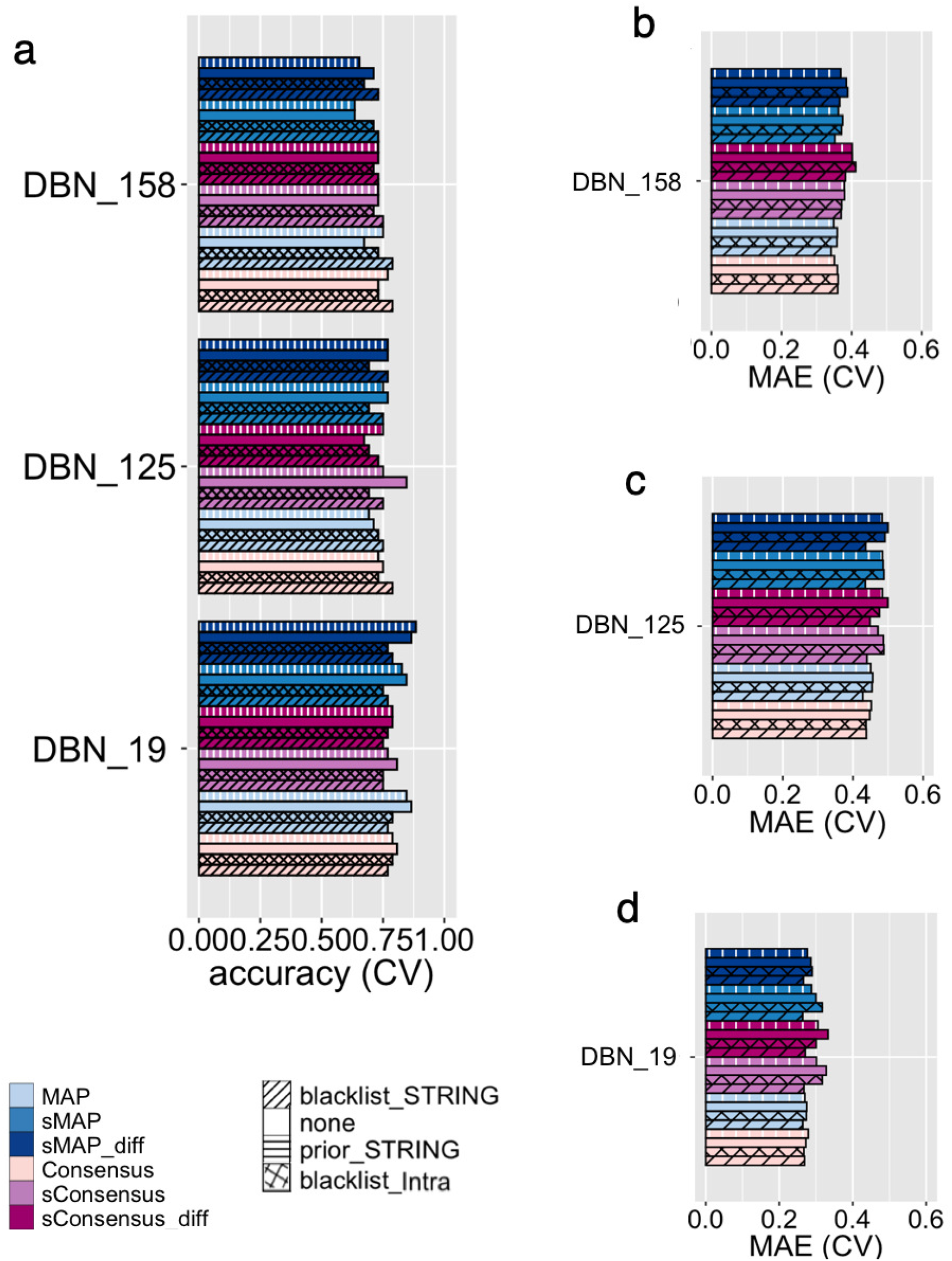
Cross-validated classification accuracy and MAE of all DBN models considered for the breast cancer dataset. DBN_19, DBN_125 and DBN_158 denote gene sets which were used to learn DBN models. Model specifications corresponding to models IDs (fill color) can be found in Table S1

**Fig. S2.**
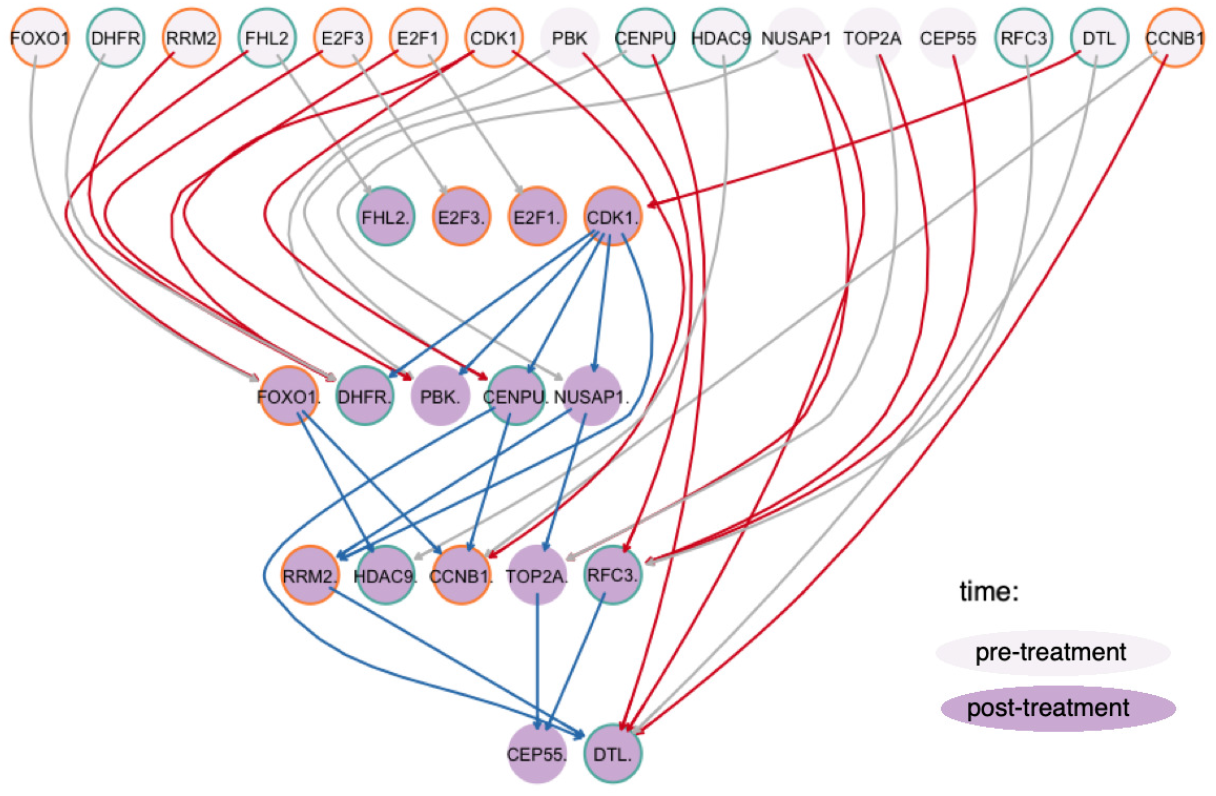
Subnetwork of the DBN transition structure estimated for the breast cancer dataset. The consensus DBN model was learned from 52 biopsies from the GSE5462 dataset. The DBN topology was restricted to edges connecting genes that were identified as functional interactors in the STRING database. The visualized subnetwork consists of genes identified as differentially expressed between responders and non-responders (green border) and their parent nodes in the estimated DBN model, including genes from p53 signaling, cellular senescence, and cell cycle pathways (orange border). Grey edges represent edges between the same genes in neighboring time slices. Red edges correspond to inter-edges. Blue edges correspond to intra-edges.

**Fig. S3.**
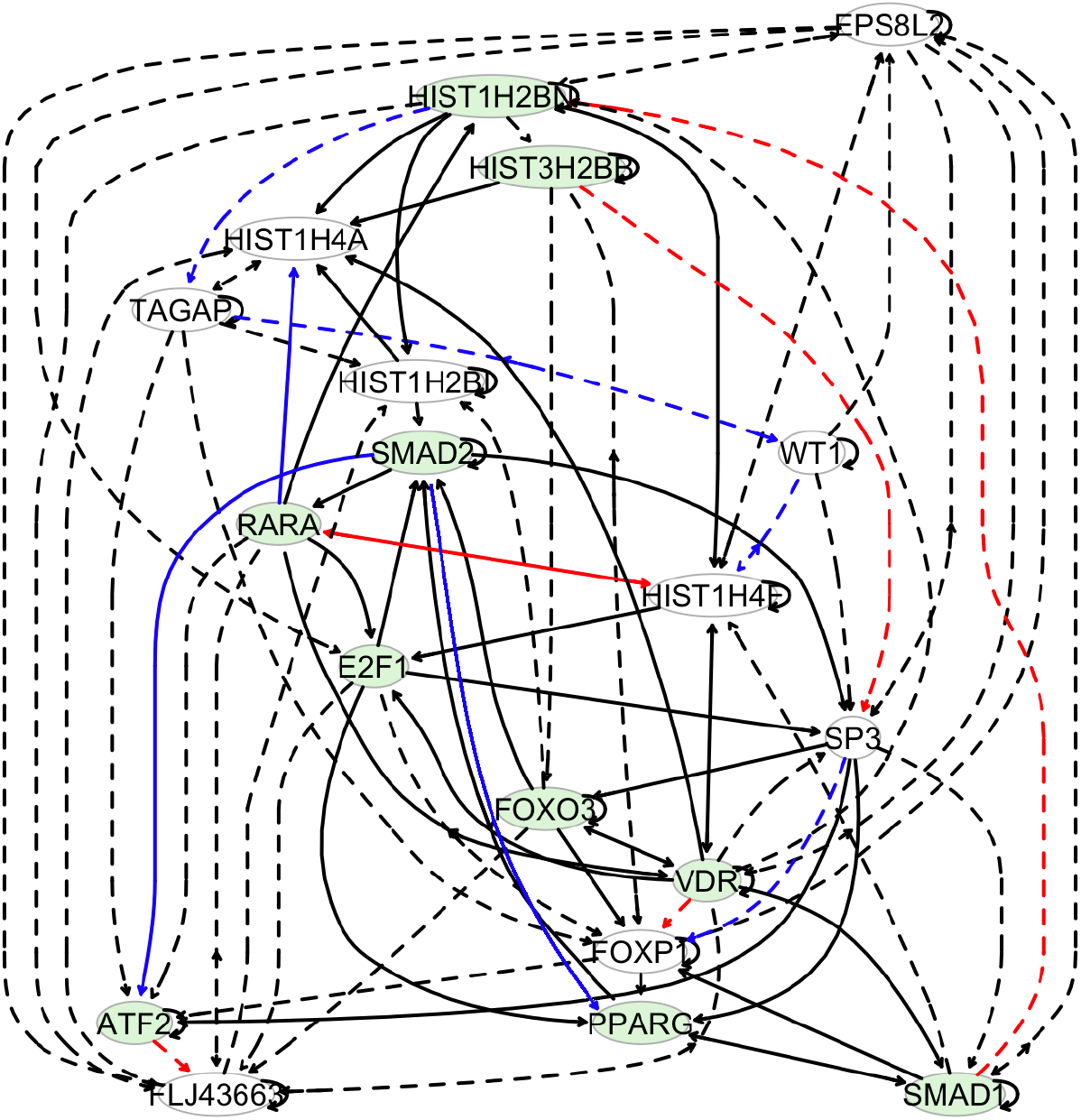
Joint subnetwork of DBN transition structures that were learned for normal and cancer samples. Intra-slice edges were blacklisted; all depicted edges are the edges between neighboring time slices. Black edges correspond to edges that are the same in two DBN models representing different subgroups. Blue edges are specific to subgroup ‘‘normal” and red edges are specific to subgroup “tumor”. Solid edges correspond to edges that were also found in the STRING database. Green nodes represent genes from the generic transcription pathway which is presented in the main text.

**Table S1.**
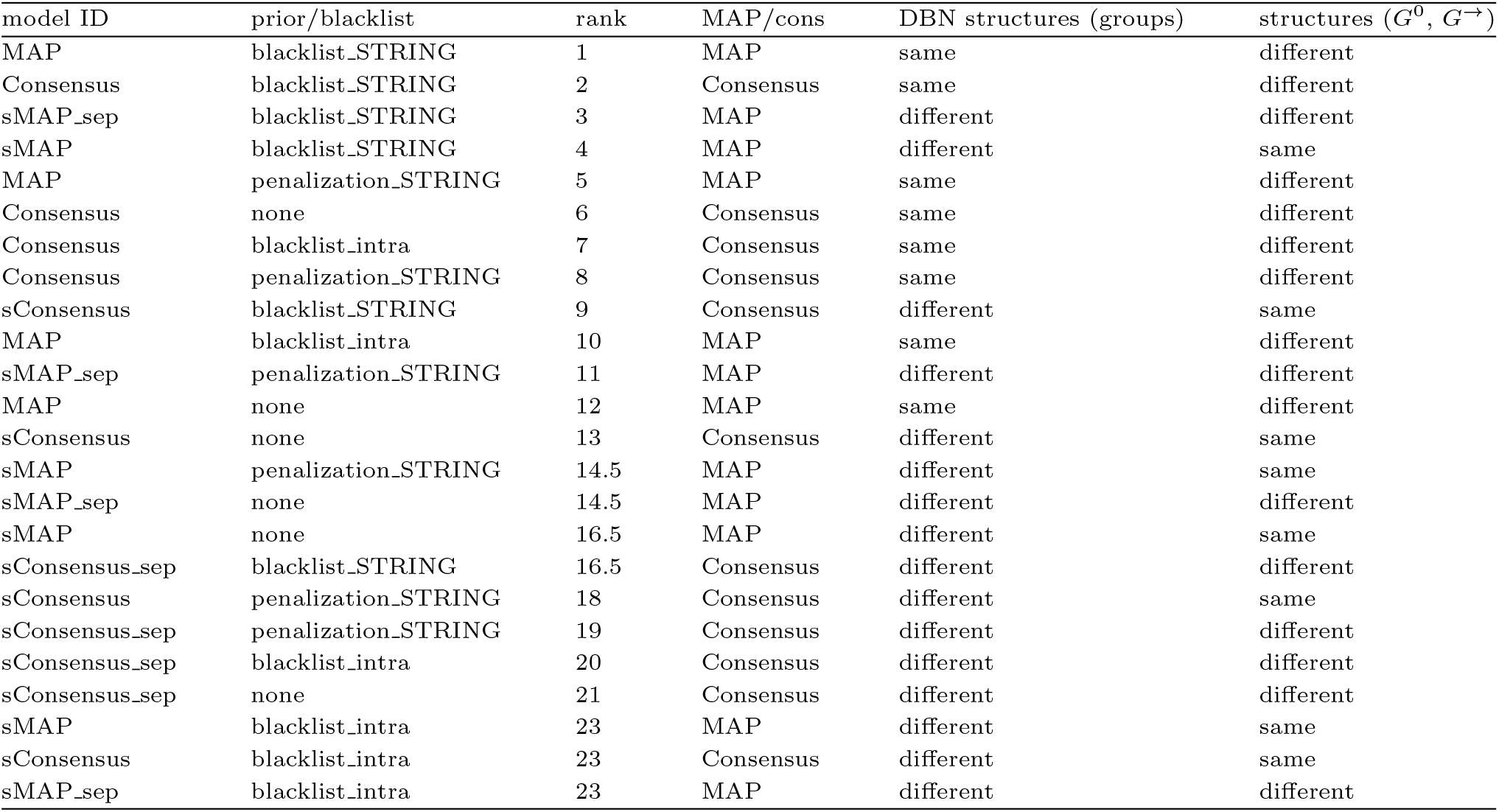
Ranking of DBN models for breast cancer dataset. To identify which DBN model performs consistently better than others for networks of different sizes and incorporate both accuracy and MAE we created a global ranking by summing the ranks of each model with regard to classification accuracy and MAE for each DBN size. Rankings in individual categories are presented in Table S2.

**Table S2.** Ranking of DBN models for breast cancer dataset. To identify which DBN model performs consistently better than others for networks of different sizes and incorporate both accuracy and MAE we created a ranking of models model with regard to classification accuracy and MAE for each DBN size; ties were resolved by taking maximum. Ranks of models in for each category are reported in this table.

| model ID | prior/blacklist | acc19 | acc125 | acc158 | mae19 | mae125 | mae158 | sum |
| --- | --- | --- | --- | --- | --- | --- | --- | --- |
| MAP | blacklist.STRING | 17.5 | 10 | 1.5 | 2 | 1 | 1 | 33 |
| Consensus | blacklist.STRING | 17.5 | 2 | 1.5 | 6 | 5 | 8 | 40 |
| sMAP_sep | blacklist.STRING | 11.5 | 4.5 | 10.5 | 3 | 3 | 11 | 44 |
| sMAP | blacklist.STRING | 17.5 | 10 | 10.5 | 1 | 2 | 4 | 45 |
| MAP | penalization.STRING | 4.5 | 21 | 4.5 | 7 | 9 | 2 | 48 |
| Consensus | none | 7.5 | 10 | 10.5 | 9 | 7 | 7 | 51 |
| Consensus | blacklist.intra | 11.5 | 15.5 | 10.5 | 4 | 4 | 9 | 54 |
| Consensus | penalization.STRING | 11.5 | 15.5 | 3 | 13 | 10 | 3 | 56 |
| sConsensus | blacklist.STRING | 22.5 | 10 | 4.5 | 5 | 6 | 13 | 61 |
| MAP | blacklist.intra | 11.5 | 15.5 | 10.5 | 10 | 11 | 5 | 64 |
| sMAP_sep | penalization.STRING | 1 | 4.5 | 22 | 12 | 15 | 12 | 66 |
| MAP | none | 2.5 | 18 | 20.5 | 11 | 12 | 6 | 70 |
| sConsensus | none | 7.5 | 1 | 10.5 | 23 | 19 | 18 | 79 |
| sMAP | penalization.STRING | 6 | 10 | 23.5 | 15 | 17 | 10 | 82 |
| sMAP_sep | none | 2.5 | 4.5 | 17.5 | 14 | 24 | 20 | 82 |
| sMAP | none | 4.5 | 4.5 | 23.5 | 17 | 18 | 16 | 84 |
| sConsensus_sep | blacklist.STRING | 22.5 | 15.5 | 10.5 | 8 | 8 | 19 | 84 |
| sConsensus | penalization.STRING | 17.5 | 10 | 10.5 | 19 | 13 | 17 | 87 |
| sConsensus_sep | penalization.STRING | 11.5 | 10 | 10.5 | 20 | 16 | 22 | 90 |
| sConsensus_sep | blacklist.intra | 17.5 | 21 | 17.5 | 18 | 14 | 24 | 112 |
| sConsensus_sep | none | 11.5 | 24 | 10.5 | 24 | 23 | 23 | 116 |
| sMAP | blacklist.intra | 22.5 | 21 | 17.5 | 21.5 | 21 | 14 | 118 |
| sConsensus | blacklist.intra | 22.5 | 21 | 17.5 | 21.5 | 20 | 15 | 118 |
| sMAP_sep | blacklist.intra | 17.5 | 21 | 20.5 | 16 | 22 | 21 | 118 |

